# Reduced temporal and spatial stability of neural activity patterns predict cognitive control deficits in children with ADHD

**DOI:** 10.1101/2024.05.29.596493

**Authors:** Zhiyao Gao, Katherine Duberg, Stacie L Warren, Li Zheng, Stephen P. Hinshaw, Vinod Menon, Weidong Cai

## Abstract

This study explores the neural underpinnings of cognitive control deficits in ADHD, focusing on overlooked aspects of trial-level variability of neural coding. We employed a novel computational approach to neural decoding on a single-trial basis alongside a cued stop-signal task which allowed us to distinctly probe both proactive and reactive cognitive control. Typically developing (TD) children exhibited stable neural response patterns for efficient proactive and reactive dual control mechanisms. However, neural coding was compromised in children with ADHD. Children with ADHD showed increased temporal variability and diminished spatial stability in neural responses in salience and frontal-parietal network regions, indicating disrupted neural coding during both proactive and reactive control. Moreover, this variability correlated with fluctuating task performance and with more severe symptoms of ADHD. These findings underscore the significance of modeling single-trial variability and representational similarity in understanding distinct components of cognitive control in ADHD, highlighting new perspectives on neurocognitive dysfunction in psychiatric disorders.

## Introduction

Childhood attention-deficit/hyperactivity disorder (ADHD) is one of the most prevalent neurodevelopmental disorders, affecting 5-8% of children worldwide ^1,2^. Hallmarks of ADHD include hyperactivity, impulsivity, and deficits in sustaining attention and behavioral control ^3,4^. Inhibitory dysregulation, the impaired ability to suppress context-inappropriate responses, is hypothesized to be a core feature of these behavioral phenotypes ^3,5–7^. Theories posit that temporal fluctuations in inhibitory control processes are central to inhibitory dysregulation in ADHD, although a crucial yet overlooked aspect is the stability and variability of neural coding during cognitive control ^8^. Distinguishing temporal fluctuations in brain responses and associated cognitive processes with underlying inhibitory control is critical for elucidating neural mechanisms and pathways characterizing the disorder’s core phenotypic features. Yet, most prior studies have largely focused on average neural responses, overlooking spatiotemporal dynamics that may provide more sensitive biological signatures of ADHD ^9,10^.

Inhibitory regulation involves two distinct processes - proactive and reactive ^11,12^. Proactive control refers to preparation of strategic responses in advance of actions, when increased control is anticipated; whereas reactive control involves inhibiting prepotent responses when interference occurs ^11–14^. Elucidating the precise nature of dysfunction associated with proactive and reactive processes is crucial for models of cognitive control deficits in ADHD pathology.

However, few studies have systematically examined the neural mechanisms underlying temporal fluctuations in proactive and reactive control and their relation to behavioral fluctuations and clinical symptoms in children with ADHD. Understanding these distinct control mechanisms and their dynamic neural representations is necessary for tailoring interventions that address specific deficits, potentially improving treatment outcomes by targeting underlying cognitive processes more accurately.

Indeed, deficits in these separate but interacting processes may underlie distinct cognitive and behavioral symptoms of ADHD ^15^. For example, impaired proactive control may contribute to problems with sustained attention, yet reactive control deficits may drive increased impulsivity ^12,16^. Disentangling the independent contributions of proactive and reactive control dysfunction to the clinical profile of ADHD will enable understanding of underlying neural mechanisms and the development of tailored interventions. Furthermore, exploring both processes should provide insight into whether ADHD involves an overall inhibitory control impairment or selective deficits in reactive stopping versus proactive preparation ^15^.

Reactive control has been extensively investigated using the Stop-Signal Task (SST), in which participants make responses to frequent go signals but must withhold their responses when infrequent stop signals occur ^17^. The stop-signal reaction time (SSRT), which estimates how fast one can cancel a prepotent response, has been widely used as an index of reactive control ^17,18^, characterizing inhibitory control deficits in clinical populations ^5,7^. Reactive control engages a widely distributed set of cortical regions, including the anterior insula (AI) and dorsomedial prefrontal cortex (dmPFC) nodes of the salience network (SN), and the dorsolateral PFC (dlPFC) and posterior parietal cortex nodes of the frontoparietal network (FPN) ^19–26^. Previous studies have shown that the AI is involved in the initiation of control processes and the dmPFC in maintaining and adjusting these processes over time ^27–30^. Disturbance or dysfunction in these regions has been linked to difficulties in maintaining sustained attention and regulating responses ^31–35^. The dlPFC and PPC are key components of the frontoparietal network, which supports executive functions such as maintenance and manipulation of information in working memory and task switching ^36–41^. These regions have been consistently implicated in studies of inhibitory control ^19,20,27^. The dlPFC is particularly important for implementing control strategies, while the PPC is crucial for integrating sensory and motor information to guide behavior ^36,42^. However, few studies have examined the temporal variability in neural coding and the spatial stability of neural representations, which limits our understanding of how ADHD impacts the brain’s ability to consistently engage in reactive control.

These challenges are further compounded when considering proactive control, an area where even less is known about the underlying neural mechanisms and their disruption in ADHD. Proactive control typically involves scenarios in which participants receive prior cues about impending tasks requiring heightened cognitive control. This approach assesses the ability to strategize and prepare for control demands in advance ^14,43–48^. Research reveals that individuals showing greater degrees of proactive control (or response slowing) also exhibit better reactive control (i.e., faster SSRT), and this is demonstrated in both adults ^43^ and children ^15^. These findings align with prior investigations demonstrating that proactive and reactive inhibitory control engage overlapping brain areas in the SN and FPN ^43–46,49–51^. Despite the recognized importance of distinguishing between proactive and reactive control in ADHD, no research has yet systematically investigated if these control processes share common neural substrates or how their potential neural coding disruptions manifest within ADHD.

As noted, a critical unexplored aspect of ADHD is neural variability and its relation to fluctuations in behavior. A prominent feature of ADHD is high intra-individual variability in behavior and symptom profiles over time ^8^. Previous behavioral studies of ADHD reveal that high intra-individual response variability (IIRV) is a consistent behavioral phenotype of ADHD ^7,10,52,53^. Attentional lapses are attributed to elevated IIRV ^54,55^, although empirical evidence demonstrating strong associations between IIRV and clinical measures of inattention is still lacking. Examining stability versus variability of task-evoked responses at the single-trial level can provide a window into the neural dynamics that may directly relate to behavioral regulation. For example, trial-level instability in activation patterns within prefrontal regions critical for cognitive control could manifest as intermittent attentional lapses or impulse control failures.

Capturing moment-to-moment fluctuations in brain function has the potential to shed light on neurobiological bases of the heterogeneity and inconsistency of ADHD symptoms.

Here we address crucial knowledge gaps in our understanding of the neural mechanisms of proactive and reactive control, through an innovative experimental design and single-trial analysis tailored for disentangling neural coding of proactive vs. reactive control. We used a cued stop-signal task (CSST) (**Figure 1a**), which allows us to directly probe neural responses associated with proactive and reactive control. Leveraging ultra-fast fMRI, we model trial-evoked responses (**Figure 1b**) to assess variability over time and similarity of spatial patterns across trials using representational similarity analysis (RSA) (**Figures 1c-d**). RSA captures fine-grained patterns, revealing whether brain regions encode information similarly across task events ^56^.

**Figure 1.**
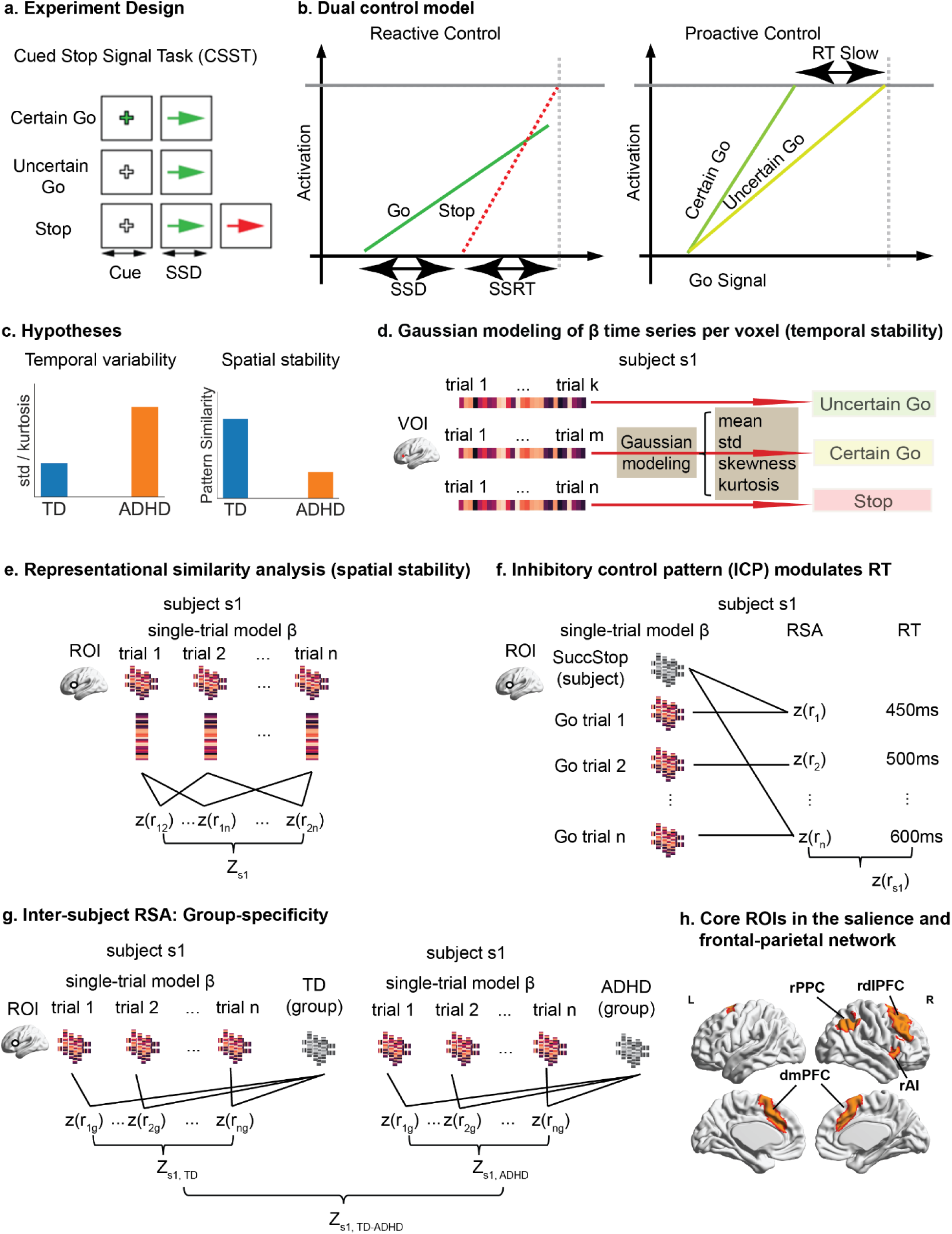
Experimental design, analysis pipeline, and hypotheses. **a.** CSST paradigm involved Certain Go, Uncertain Go and Stop trials. **b.** Dual control model decomposed reactive control and proactive control processes in the CSST. **c.** We hypothesized that children with ADHD will show higher temporal variability and lower spatial stability of trial-evoked brain responses than TD children. **d.** The Gaussian model was applied to assess temporal variability of trial-evoked neural responses. **e.** Representational similarity analysis (RSA) was used to examine spatial stability of trial-evoked activation patterns. **f.** ICP index was computed to measure the extent to which proactive control was implemented in Certain and Uncertain Go trials, which modulates trial-wise behavioral fluctuation. **g.** Inter-subject pattern similarity analysis was used to investigate group-specificity of neural coding during proactive and reactive control. **h.** Core regions of interests (ROIs) in the salience and frontal-parietal network, defined from an independent meta-analysis study (Shirer et al., 2012). CSST, stop-signal task; SSD, stop-signal delay; SSRT: stop-signal reaction time; ICP: inhibitory control pattern; rAI, right anterior insula; dmPFC, dorsal medial prefrontal cortex; rdlPFC, right dorsal lateral prefrontal cortex; rPPC, right posterior parietal cortex; RT: reaction time; RSA: representational similarity analysis.

This approach goes beyond previous research that has relied on average activation levels, enabling us to capture temporal variations in neural representations across trials. Examining inter-trial variability and consistency of neural coding allows us to relate neural instability directly to the heterogeneity and fluctuation of behavioral regulation in children with ADHD. This novel experimental and analytic approach elucidates whether and how neural network dynamics differ in children with ADHD versus TD children.

We hypothesized that children with ADHD would show increased temporal variability and reduced spatial stability of trial-evoked responses in the AI, dmPFC, dlPFC and PPC nodes of the SN and FPN; weakened association between neural coding of inhibitory control and fluctuating behavior; decreased within-group similarity of trial-evoked responses in the SN and FPN; and temporal and spatial variability of trial-evoked response in association with clinical symptoms. Our findings of neural instability advance models of inhibitory control dysfunction by providing insights into the neural mechanisms that instantiate the heterogeneous behavioral symptoms of ADHD. More generally, advanced neuroimaging techniques allowed us to move beyond traditional methods that average neural activity over trials, providing a more detailed and accurate picture of how neural variability contributes to cognitive control deficits in ADHD.

## Results

### Weak and unstable dual control in children with ADHD

We recruited 107 children (9-12 years old) from the local community, comprising 53 children with ADHD and 40 TD children, who all completed two runs of the CSST. The final fMRI-sample included 26 children with ADHD (10.8±1.2 years old, 10F/16M) and 35 TD children (10.5±1.1 years old, 13F/22M) who met task performance and head motion criteria (see the Method section for details). Both groups were matched in age and gender (all *ps*>0.2, two-sample t-test, **Table 1**). In comparison to TD children, children with ADHD had significantly higher inattention and hyperactivity/impulsivity scores (all *ps* < 0.001), greater head motion and lower full-scale IQ (all *ps*<0.05, two-sample t-test, **Table 1**).

**Table 1.**
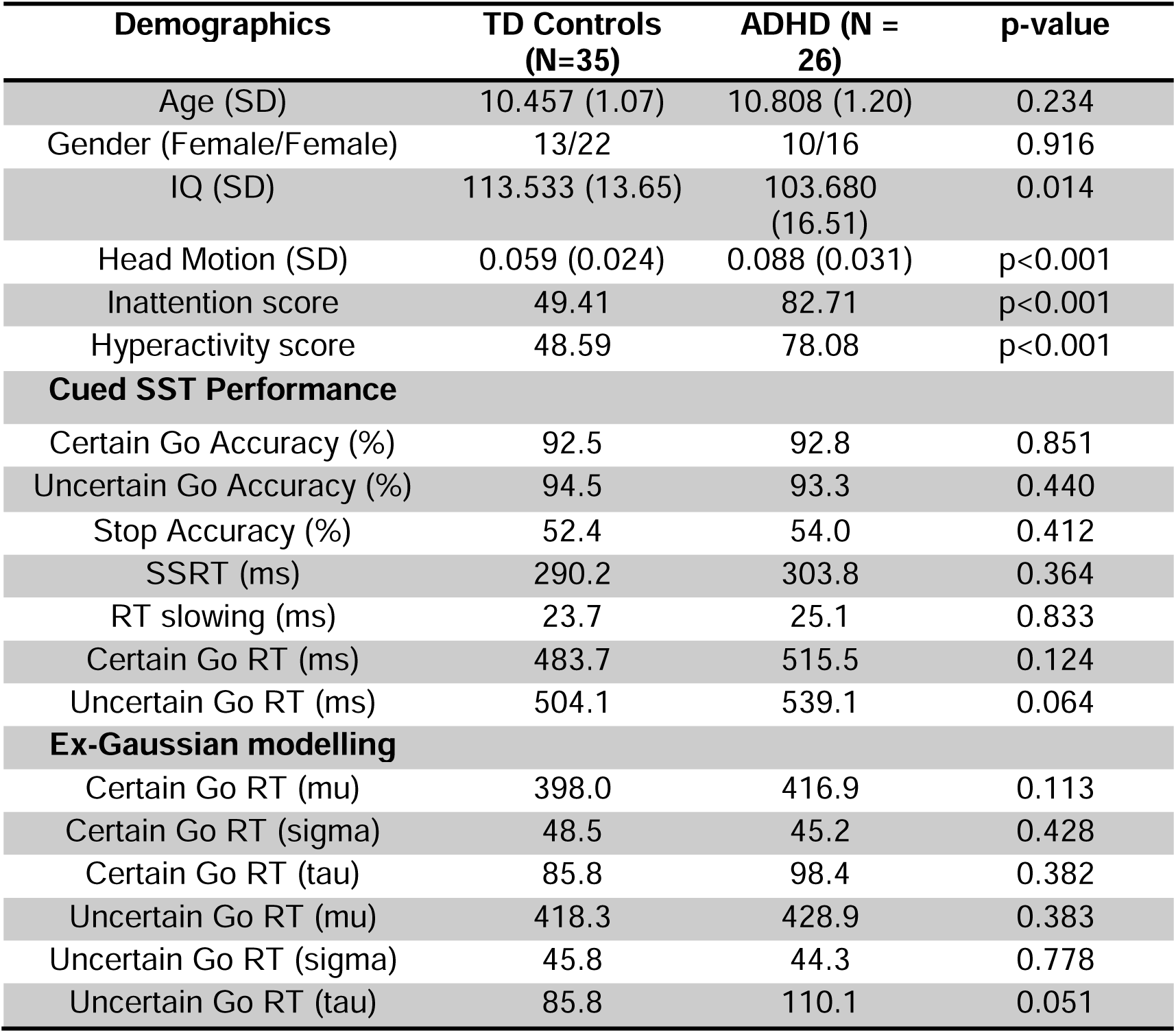
Demographics and behavioral results (fMRI sample)

| **Demographics** | **TD Controls**<br>**(N=35)** | **ADHD (N =**<br>**26)** | **p-value** |
| --- | --- | --- | --- |
| Age (SD) | 10.457 (1.07) | 10.808 (1.20) | 0.234 |
| Gender (Female/Female) | 13/22 | 10/16 | 0.916 |
| IQ (SD) | 113.533 (13.65) | 103.680<br>(16.51) | 0.014 |
| Head Motion (SD) | 0.059 (0.024) | 0.088 (0.031) | p<0.001 |
| Inattention score | 49.41 | 82.71 | p<0.001 |
| Hyperactivity score | 48.59 | 78.08 | p<0.001 |
| **Cued SST Performance** |  |  |  |
| Certain Go Accuracy (%) | 92.5 | 92.8 | 0.851 |
| Uncertain Go Accuracy (%) | 94.5 | 93.3 | 0.440 |
| Stop Accuracy (%) | 52.4 | 54.0 | 0.412 |
| SSRT (ms) | 290.2 | 303.8 | 0.364 |
| RT slowing (ms) | 23.7 | 25.1 | 0.833 |
| Certain Go RT (ms) | 483.7 | 515.5 | 0.124 |
| Uncertain Go RT (ms) | 504.1 | 539.1 | 0.064 |
| **Ex-Gaussian modelling** |  |  |  |
| Certain Go RT ($\mu$) | 398.0 | 416.9 | 0.113 |
| Certain Go RT ($\sigma$) | 48.5 | 45.2 | 0.428 |
| Certain Go RT ($\tau$) | 85.8 | 98.4 | 0.382 |
| Uncertain Go RT ($\mu$) | 418.3 | 428.9 | 0.383 |
| Uncertain Go RT ($\sigma$) | 45.8 | 44.3 | 0.778 |
| Uncertain Go RT ($\tau$) | 85.8 | 110.1 | 0.051 |

Because participants who have more severe clinical symptoms of ADHD are more likely to be excluded, for example for excessive head motion, we conducted a subsequent sensitivity analysis including participants who have large head motion in the behavioral analysis. The larger behavior-only sample includes 50 children with ADHD and 37 TD children. Both groups were matched in age and gender (all *ps*>0.4, **Supplementary Table S1**).

Reactive control was estimated using the SSRT. We did not find a significant group difference in SSRT in the fMRI sample (*t*_59_=1.13, *p*=0.26, 95% CI=[-0.049, 0.013], Cohen’s d=0.293). Yet, in the behavior-only sample, we found that children with ADHD had significantly longer SSRT than TD children (*t_85_*=2.46, *p*=0.016, 95% CI=[0.008, 0.074], Cohen’s d=0.532, two sample t-tests).

Proactive control was estimated by the RT difference between Uncertain and Certain Go trials. Both children with ADHD and TD children exhibited significant proactive control, i.e. response slowing in Uncertain versus Certain Go trials (*ps*<0.01, single-sample two-tailed t-test). But there was no significant group difference in the fMRI and behavior-only samples (*ps*>0.7).

We also examined IIRV, quantified by the *sigma* and *tau* of the ex-Gaussian model ^57^, in the Certain and Uncertain Go trials separately. There were no significant group differences in sigma and tau in either trial type in the fMRI sample (all *ps*>0.05). However, in the behavior-only sample, children with ADHD showed significantly higher tau than TD children in Uncertain (*t_85_*=3.23, *p*=0.002, 95% CI = [0.013, 0.052], Cohen’s d = 0.70) and Certain Go trials (*t_85_*=2.36, *p*=0.02, 95% CI = [0.038, 0.451], Cohen’s d = 0.51).

Together, these behavioral findings suggest that, in comparison to their TD peers, children with ADHD have poor reactive control and less stable performance but preserved proactive control functions.

### Increased temporal variability of trial-evoked brain activation in children with ADHD

Next, we assessed the temporal variability of trial-by-trial brain activation during the CSST. First, we conducted a single-trial GLM analysis to estimate the trial-evoked brain response for each single trial. Then, we fit the Gaussian model to the trial-wise activation per voxel per task condition, from which the four model parameters i.e., mean, standard deviation (STD), skewness, and kurtosis were calculated. The normal distribution of trial-evoked brain responses was assessed (**Supplementary Methods and Supplementary Table S2**). Significance of between-group comparison was determined using the threshold-free cluster enhancement method (TFCE, *p*<0.05 corrected).

First, no significant between-group difference in mean was found in any task condition.

Then, we examined STD of trial-evoked neural responses. In comparison to TD children, children with ADHD exhibited significantly higher STD in the frontal, temporal and cingulate regions, including the bilateral inferior frontal gyrus, dmPFC, precuneus, and lateral and medial temporal lobe, in Uncertain and Certain Go and Stop trials (**Figure 2a)**.

**Figure 2.**
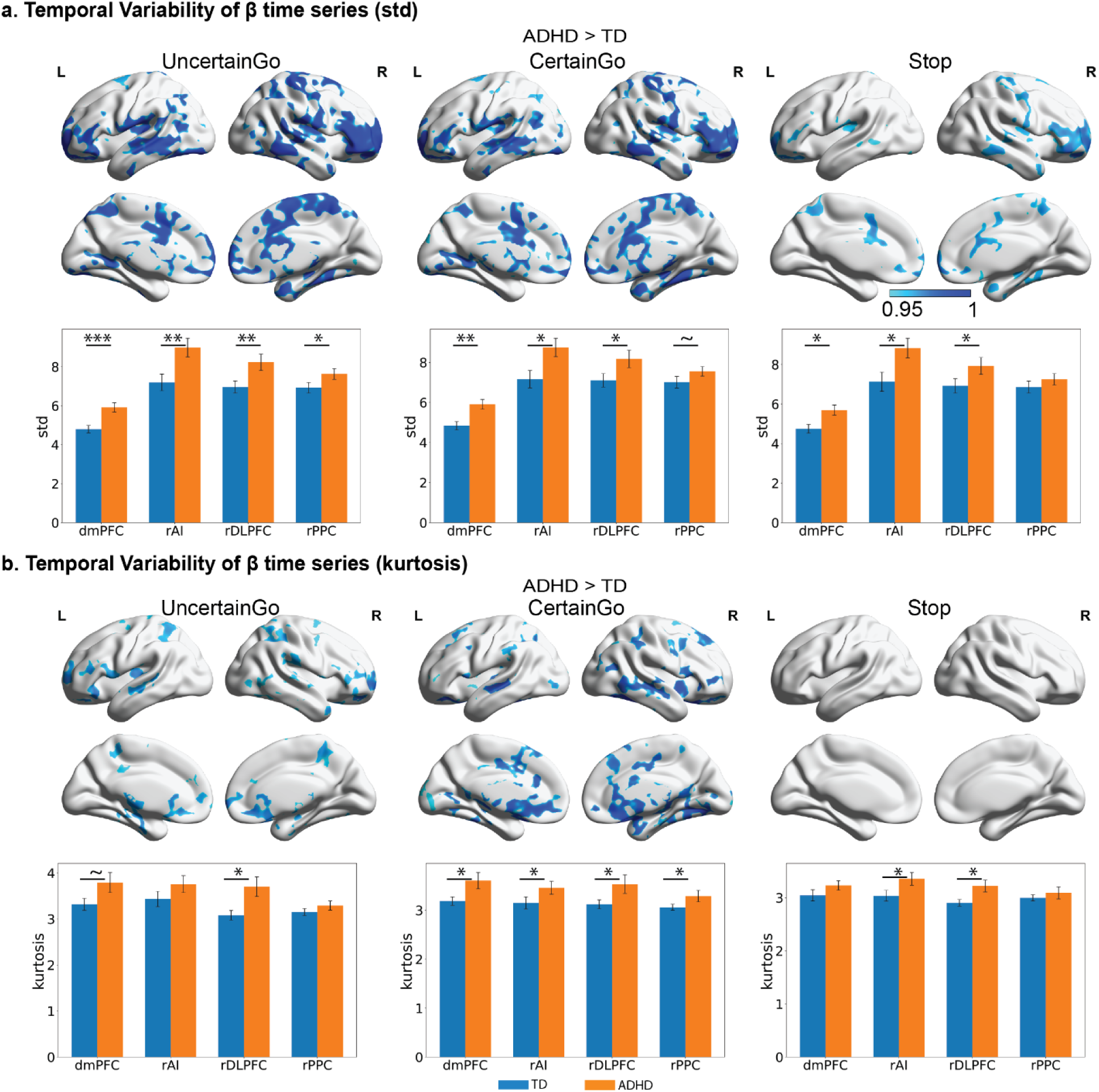
**Heightened temporal variability of neural responses in children with ADHD**. **a.** Children with ADHD showed significantly greater variability, measured using standard deviation (std), than typically developing (TD) controls (*p*<0.05, corrected). **b.** Children with ADHD also showed significantly greater kurtosis than TD children (*p*<0.05, corrected). std: standard deviation.

Next, we measured kurtosis of trial-evoked neural responses. Kurtosis measures the degree of tailedness in a data distribution. Higher kurtosis is often associated with heavy tails or more outliers. In comparison to TD children, children with ADHD had significantly higher kurtosis in the lateral and ventromedial prefrontal, posterior insular, posterior parietal, and temporal regions in Uncertain and Certain Go trials (**Figure 2b)**.

Last, we examined skewness of trial-evoked neural responses. Skewness measures the degree of asymmetry in a data distribution. Positive skew indicates that data distribution has a long right tail (i.e. greater activation) whereas negative skew indicates that data distribution has a long left tail (i.e. weaker activation or deactivation). Children with ADHD showed significantly smaller skewness than TD children in the frontal, parietal and cingulate regions, including bilateral inferior frontal gyrus, middle frontal gyrus, superior frontal gyrus, dmPFC, posterior cingulate cortex/precuneus, and PPC, in Uncertain Go trials (**Supplementary Figure S1)**.

Taken together, children with ADHD demonstrated greater temporal variability of trial-evoked neural responses (larger STD) with more outlier activation (greater kurtosis) in the SN and FPN, suggesting unstable recruitment of core cognitive control systems during task performance.

Note that we also conducted control analyses using conventional GLM to examine averaged task-evoked brain activation but did not find significant between-group difference in any task conditions.

To test the robustness of our findings, we conducted additional ROI-based analyses focusing on standard deviation and kurtosis using independent ROIs implicated in cognitive control from salience (SN), encompassing the right anterior insula (rAI) and dorsal medial prefrontal cortex (dmPFC), and frontal-parietal regions (FPN), encompassing right dorsal lateral prefrontal cortex (rdlPFC), and right posterior parietal cortex (rPPC) (see Methods and **Figure 1h**). In comparison to TD children, children with ADHD exhibited significantly higher standard deviation in all four ROIs during Uncertain Go trials, in the rAI, dmPFC and rdlPFC during Certain Go trials, and Stop trials (all ps<0.05, FDR corrected, **Figure 2a**). Moreover, children with ADHD had significantly higher kurtosis than TD children in the rdlPFC during Uncertain Go trials, the rAI, rdlPFC, and rPPC during Certain Go trials, and the rAI and rdlPFC during Stop trials (all *ps*<0.05, **Figure 2b**).

### Temporal variability of trial-evoked brain responses in relation to core symptoms of ADHD

We also examined whether temporal variability of trial-evoked brain responses is associated with the core clinical symptoms of ADHD focusing on the key ROIs implicated in cognitive control. For brain measures, we used kurtosis because kurtosis is particularly influenced by extreme values. After controlling the age, gender, IQ, and head motion, inattention scores were marginally significantly correlated with the kurtosis in the right PPC (*r*_partial_=0.33, *p*=0.074, FDR corrected) during Certain Go trial, suggesting that higher kurtosis is associated with more severe inattention problems. We further examined the relationship within each group and our results found that only children with ADHD demonstrated a significant positive relationship between the kurtosis in the rPPC during Certain Go trials and inattention symptoms after controlling confounding factors (*r*_partial_=0.573, *p*=0.033, FDR corrected). These findings suggest that temporal variability of trial-evoked brain response has clinical implications.

### Weak spatial stability of trial-evoked brain response patterns in children with ADHD

Next, we examined whether spatial stability of trial-evoked brain responses during dynamic proactive and reactive control is weakened by the disorder. Spatial stability was quantified by the similarity (i.e., *Pearson’s* correlation) of voxel-wise activation patterns between any pair of the same type of trials and a searchlight approach was implemented to produce a whole brain map of task-dependent spatial stability per subject ^56,58^. Significance of spatial stability was determined using the threshold-free cluster enhancement method (TFCE, *p*<0.05 corrected).

We found that children with ADHD demonstrated significantly weaker spatial stability in bilateral inferior frontal gyrus, dmPFC, the lateral parietal cortex, and ventral/medial temporal lobe in both Certain and Uncertain Go trials than TD children (*p*<0.05, corrected, **Figure 3a**).

**Figure 3.**
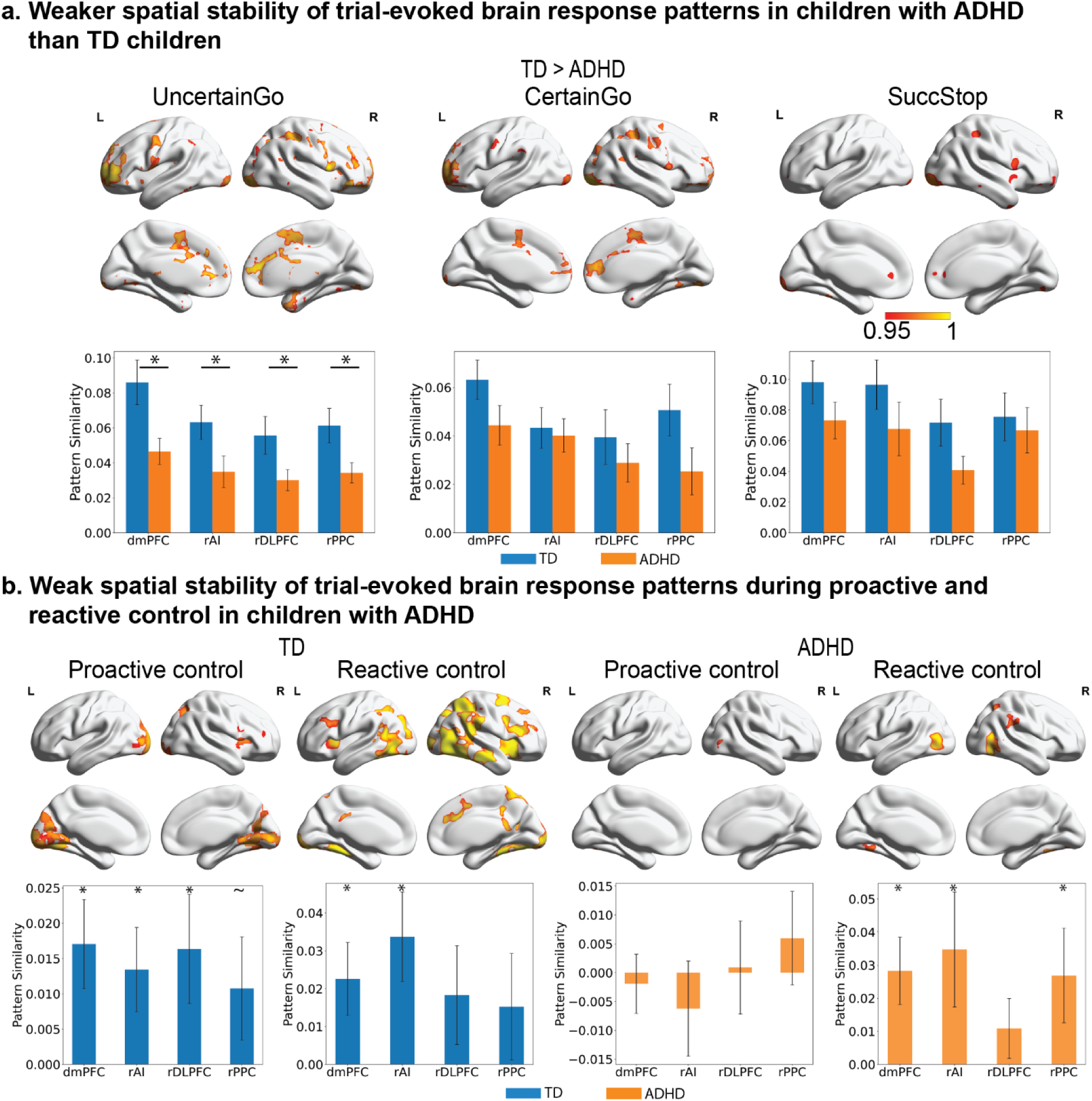
Weakened spatial stability of trial-evoked neural responses in children with ADHD. **a.** Typically developing (TD) children showed greater spatial stability than children with ADHD in salience and frontoparietal networks during Uncertain Go, Certain Go and SuccStop trials. **b.** Highly stable recruitment of salience and frontoparietal networks during proactive and reactive control was found in TD children (*p*<0.05, corrected) but not in children with ADHD.

Additionally, children with ADHD have lower spatial stability in the bilateral AI during Uncertain Go trials than TD children (*p*<0.05, corrected, **Figure 3a**). Children with ADHD also exhibited reduced spatial stability in the right inferior frontal gyrus, AI, PPC, and dmPFC during successful Stop (SuccStop) trials in children compared to TD children (*p*<0.05, corrected, **Figure 3a**).

We did not find significantly increased spatial stability in any brain region in children with ADHD in comparison to TD children.

ROI-based analyses revealed that, in comparison to TD children, children with ADHD demonstrated significantly lower spatial stability in all four ROIs during Uncertain Go trials (all *ps*<0.05, FDR corrected, **Figure 3a**). However, there were no significant effects found during Certain Go and Successful Stop trials (all *ps*>0.11).

Together, these results suggest that children with ADHD recruit highly variable distributed brain regions from trial to trial during cognitive control performance in comparison to TD children.

### Weak spatial stability of trial-evoked brain response patterns during proactive and reactive control in children with ADHD

Because trials involving the same cognitive processes would exhibit greater representational similarity compared to trials that involves different processes, it allows us to identify representational brain systems uniquely associated with proactive and reactive control.

Proactive control was measured by the RSA contrast between Uncertain and Certain Go trials, and reactive control was measured by the RSA contrast between Successful Stop and Uncertain Go trials. For proactive control, TD children showed significant spatial stability in the right AI, inferior frontal gyrus, and bilateral visual cortex, whereas children with ADHD showed weak effect only in visual areas (*p*<0.05, corrected, **Figure 3b**). For reactive control, TD children showed significantly spatial stability in the bilateral AI, inferior frontal gyrus, PPC, anterior and posterior cingulate cortex and visual areas, whereas children with ADHD showed significant spatial stability in less distributed brain regions, including visual areas and right PPC (*p*<0.05, corrected, **Figure 3b**). While there was no significant between-group difference, children with ADHD showed less spatial stability in the right AI, inferior frontal gyrus and superior frontal gyrus during proactive control and the right AI, left middle frontal gyrus and middle temporal gyrus during reactive control than TD children in a less stringent threshold (*p*<0.01, uncorrected; **Supplementary Figure S2**. Together, our findings suggest that children with ADHD have weakened spatial stability in the distributed brain systems during proactive and reactive control in comparison to TD children.

Further ROI analyses revealed that TD children showed great spatial stability of trial-evoked neural responses in the rAI, dmPFC and rdlPFC during proactive control and in the rAI and dmPFC during reactive control (*p*<0.05, FDR corrected, **Figure 3b**). Children with ADHD also exhibited great spatial stability in the rAI and dmPFC (*p*<0.05, FDR corrected, **Figure 3b**) but not during proactive control (all *ps*>0.7).

### Weak association between trial-evoked inhibition-alike brain response and RT fluctuation in children with ADHD

Increased IIRV is a robust behavioral phenotype associated with ADHD, which is commonly regarded as an index of attention lapse or inattention ^10,15,59^. However, behavioral fluctuation could also be driven by trial-by-trial response strategy adjustment, a dynamic proactive control process modulated by participants’ time-varying anticipation ^15,21,60,61^. Here we examined whether the degree of proactive control in trial-evoked brain response patterns can track RT fluctuations in Certain and Uncertain Go trials. As proactive and reactive control may share similar neural underpinnings ^46^, we developed an “inhibitory control pattern” (ICP) index in each Certain and Uncertain Go trial by quantifying the similarity between trial-evoked brain response pattern on Go trial and a subject-specific “template” brain response pattern of inhibitory control (**Figure 1e**). The subject-specific “template” brain response pattern of inhibitory control was obtained by averaging trial-evoked brain response pattern from all the Successful Stop trials (β-maps from the single-trial model). Then a searchlight approach was implemented to produce a whole brain map of ICP index in Certain and Uncertain Go trials separately per subject (see Methods for more details). The larger ICP value, the greater similarity between trial-evoked brain response pattern on Certain and Uncertain Go trials and the averaged brain response pattern on Successful Stop trials, indicating greater engagement of inhibitory control systems during Go trials. Last we examined correlation between voxel-wise ICP and RT across trials.

We found that trial-wise RT fluctuation is significantly correlated with the ICP in the bilateral AI, right inferior frontal gyrus, superior frontal gyrus, dmPFC in TD children in Certain and Uncertain Go trials (*p*<0.05, corrected, **Figure 4a**). Children with ADHD showed less distributed effect in the supplementary motor area and anterior cingulate cortex in Certain and Uncertain Go trial (*p*<0.05, corrected, **Figure 4a**). Between group comparison further revealed that children with ADHD have significantly weaker relationships between trial-wise RT fluctuation and ICP in anterior cingulate cortex/supplementary motor cortex, right AI, right posterior middle temporal gyrus, and right supramarginal gyrus extending to the postcentral gyrus in Certain and Uncertain Go trials (*p*<0.05, corrected, **Figure 4b**). These findings suggest that TD children can flexibly recruit inhibitory control systems to implement proactive control and thereby modulate trial-wise response time, but such ability is much weakened in children with ADHD.

**Figure 4.**
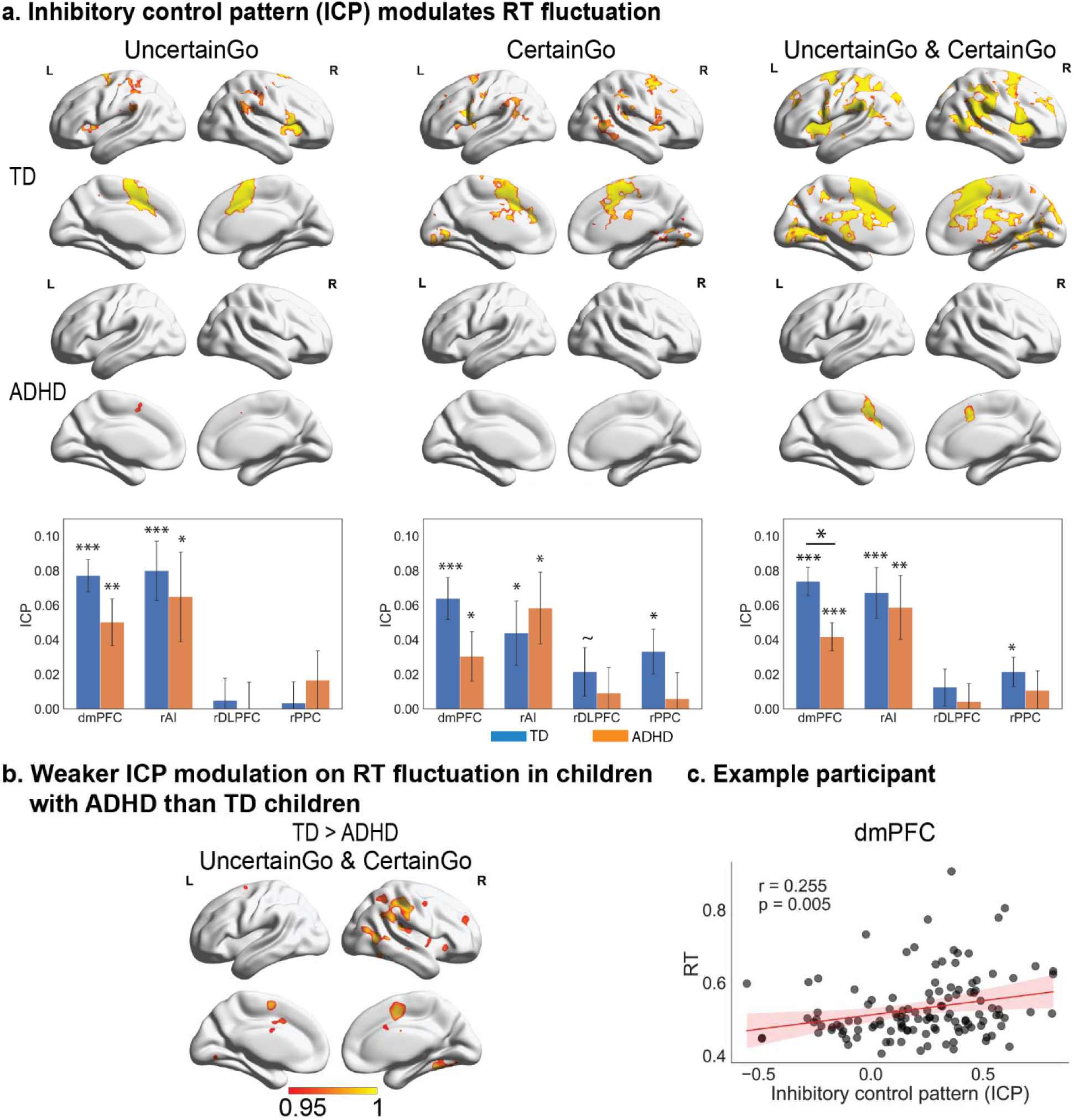
Atypical neural dynamics underlie impaired behavioral regulation in children ADHD. **a.** Inhibitory control pattern (ICP) index of the salience and frontoparietal networks during Certain and Uncertain Go trials modulates trial-wise reaction time (RT) in TD children and children with ADHD (*p*<0.05, corrected). **b.** TD children demonstrated significantly greater association between ICP index of ROIs from the salience network and trial-wise RT fluctuation than children with ADHD (*p*<0.05, corrected). **c.** Data from an exemplary participant illustrated a positive association between ICP of the dmPFC and RT. dmPFC: dorsomedial prefrontal cortex.

Additional ROI-based analyses were conducted to examine whether ICP estimated from independent ROIs were able to track the RT fluctuation for each group. TD children demonstrated a significant relation between trial-wise RT fluctuations across trial types and ICP of the rAI and dmPFC during Uncertain Go, the rAI, dmPFC and rPPC during Certain Go, and the rAI, dmPFC, and rPPC during Certain and Uncertain Go (*ps*<0.05, FDR corrected, **Figure 4a**). Children with ADHD also demonstrated significant relationships between trial-wise RT fluctuations and ICP in the rAI and dmPFC during Certain and/or Uncertain Go (*ps*<0.05, FDR corrected, **Figure 4a**), but not in the other ROIs (all *ps*>0.2). Moreover, in comparison to children with ADHD, TD children exhibited significant stronger association between trial-wise RT fluctuations and the ICP of dmPFC during both Uncertain and Certain Go trials (*p*=0.016) but not in other ROIs (all *ps*>0.3).

### Group-specific, inter-subject similarity patterns underlying proactive and reactive control

Next, we examined whether children with ADHD demonstrated weakened spatial stability of trial-evoked brain responses within their respective groups. Notably, for a participant’s within-group spatial similarity, the participant’s own brain response pattern was excluded when computing the averaged spatial pattern of brain response of the group that the participant belongs to, which is to avoid overestimation of the within-group spatial similarity. The searchlight algorithm was used to estimate within-group inter-subject similarity across the whole brain and its statistical significance was determined using the threshold-free cluster enhancement method (TFCE, *p*<0.05 corrected).

We found that, in comparison to TD children, children with ADHD demonstrated significantly weakened inter-subject spatial stability in distributed areas in bilateral inferior frontal gyrus, left AI, PPC, precentral gyrus, right middle frontal gyrus, superior frontal gyrus, dmPFC/supplementary motor cortex, bilateral middle and inferior temporal lobe, visual cortex, and subcortical regions including caudate and thalamus in Uncertain Go trials; in precentral/postcentral gyrus and left inferior frontal gyrus in Certain Go trials; and in right AI, inferior frontal gyrus, PPC, and bilateral visual cortex in SuccStop trials (*p*<0.05, corrected, **Figure 5a**). Children with ADHD also exhibited strengthened spatial stability in visual cortex across conditions; in bilateral frontal pole, and supramarginal gyrus in Uncertain Go trials; and in supplementary motor cortex, and thalamus in SuccStop trials (*p*<0.05, corrected, **Figure 5a**).

**Figure 5.**
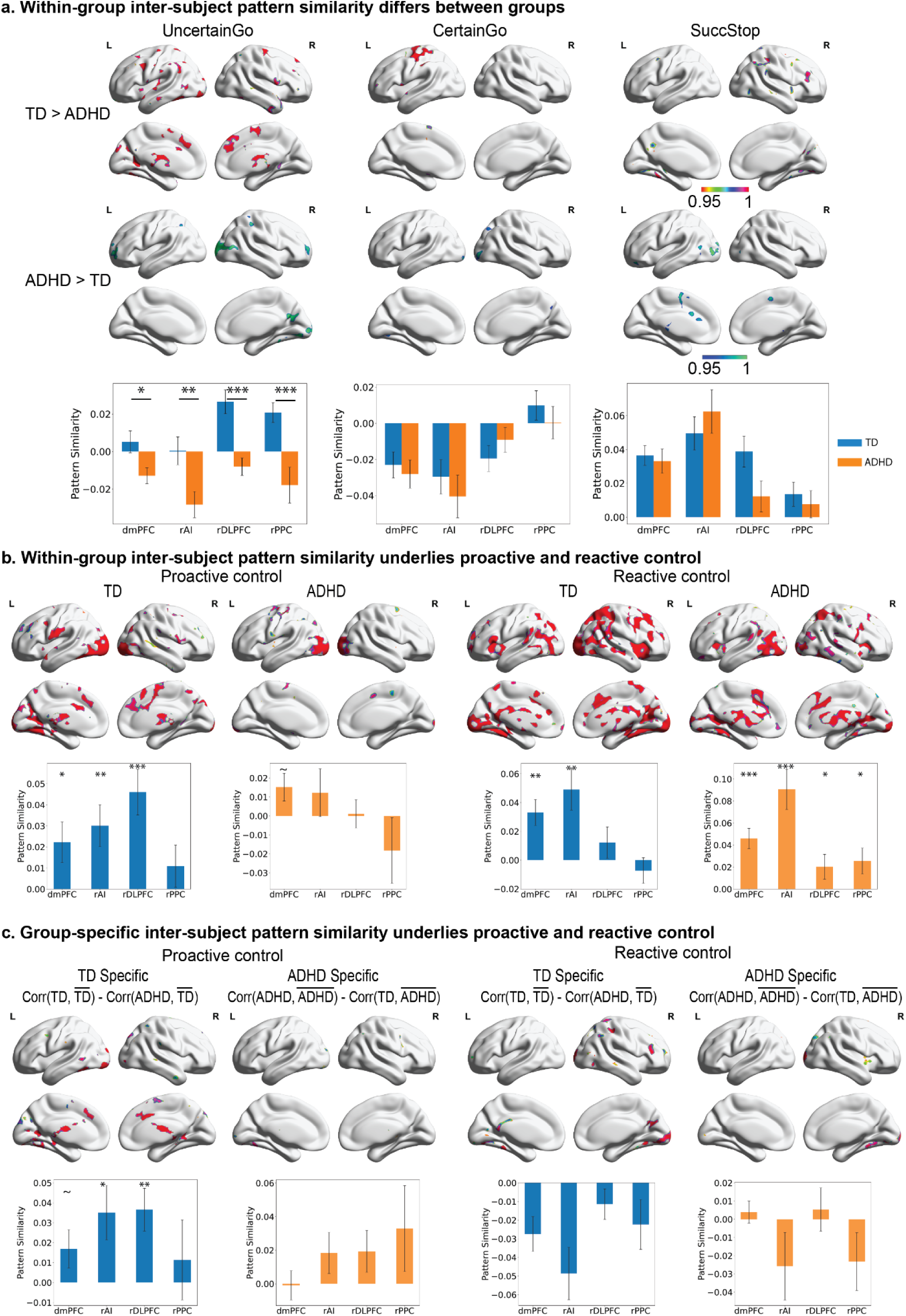
Heterogeneity of trial-evoked neural responses in children with ADHD. **a.** Within-group inter-subject spatial pattern stability analyses revealed between group differences during Certain Go, Uncertain Go and Stop trials (*p*<0.05, corrected). **b.** Within-group inter-subject spatial pattern stability during proactive and reactive control. **c.** Group-specificity of inter-subject spatial pattern similarity analyses revealed more heterogeneous trial-evoked neural response during proactive and reactive control in children with ADHD than typically developing (TD) children (*p*<0.05, corrected).

We also conducted additional ROI-based analysis to validate our findings in the key regions implicated in cognitive control. In comparison to children with ADHD, TD children demonstrated significant higher inter-subject spatial stability in all four ROIs during Uncertain Go trials (all *ps*<0.05, FDR corrected, **Figure 5a**). We did not observe significant differences in the ROIs during Certain Go and SuccStop (all *ps*>0.4).

Furthermore, we asked whether children with ADHD not only have weakened inter-subject spatial stability in each type of trials but also show more variable inter-subject spatial patterns of brain response during proactive and reactive control than TD children. So, we examined the within-group inter-subject spatial similarity of brain response during proactive and reactive control per group. The dmPFC, bilateral inferior frontal gyrus, left precentral/postcentral area, and left lateral and medial temporal lobe, and visual cortex showed great inter-subject pattern similarity during proactive control in TD children (*p*<0.05, corrected, **Figure 5b**), whereas a less distributed pattern was found in children with ADHD, including a smaller cluster in dmPFC, visual and motor cortex. The bilateral AI, dmPFC, PCC, lateral and medial temporal lobe, and basal ganglia showed great inter-subject pattern similarity during reactive control for both TD and ADHD group (p<0.05, corrected, **Figure 5b**).

ROI-based analyses found significant inter-subject spatial similarity of brain responses for TD children in the rAI, dmPFC, and rdlPFC during proactive control (*ps*<0.05, FDR corrected, **Figure 5b**), but not in the rPPC (*p*=0.14). TD children also demonstrated significant spatial stability in the rAI and dmPFC during reactive control (*ps*<0.05, FDR corrected, **Figure 5b**), but not in rdlPFC and rPPC. Children with ADHD also exhibited significant inter-subject spatial similarity of brain responses in all the ROIs during reactive control (*ps*<0.05, FDR corrected, **Figure 5b**) but not during proactive control (all ps>0.3).

Next, we examined the group specificity of inter-subject spatial similarity of brain response during proactive and reactive control per group. For each participant, we computed within-group inter-subject spatial similarity, i.e. the spatial similarity between a participant and the group of the participant, and between-group inter-subject spatial similarity, i.e. the spatial similarity between a participant and the group that the participant does not belong to (**Figure 1f**). The group specificity was defined by the difference between within-group inter-subject spatial similarity and between-group inter-subject spatial similarity. Positive values indicate that a participant’s regional activation pattern is more similar to the rest of the participants from the same group than the participants from the other group, whereas negative values indicate vice versa. During proactive control, we observed greater spatial similarity among TD children than between TD children and children with ADHD in the supplementary motor cortex, dmPFC, bilateral posterior middle temporal gyrus, bilateral medial temporal lobe, as well as subcortical clusters, including the thalamus and caudate (*p*<0.05, corrected, **Figure 5c**). During reactive control, the right inferior frontal gyrus, PCC, middle temporal gyrus, and lateral parietal cortex show greater spatial similarity among TD children than between TD children and children with ADHD during reactive control, whereas the right AI showed greater similarity among children with ADHD than TD children (p<0.05, corrected, **Figure 5c**).

ROI-based analysis was conducted to examine the group-specific inter-subject spatial similarity during proactive and reactive control in key regions of the SN and the FPN within each group. Only TD children demonstrated significant spatial stability in the rAI and rdlPFC during proactive control (*ps*<0.05, FDR corrected, **Figure 5c**) but not in the dmPFC and rPPC (*ps*>0.05). We did not find a significant group specificity during proactive control for children with ADHD (all *ps*>0.1). Additionally, there was no significant group specificity during reactive control for both children with ADHD and TD children (all *ps*>0.5).

Last, we conducted ROI-based analysis to investigate whether similarity to the averaged spatial patterns of the ADHD group during proactive and reactive control is associated with individual differences in core symptoms of ADHD. We found a significant positive correlation between the pattern similarity to ADHD group averaged spatial pattern during proactive control in rAI and inattention score (*r_partial_*=0.436, *p*<0.007, FDR corrected, **Figure 6**) after accounting for age, gender, IQ, and head motion. Our results indicated that a more ADHD-like neural activity pattern during proactive control was associated with more severe clinical symptoms.

**Figure 6.**
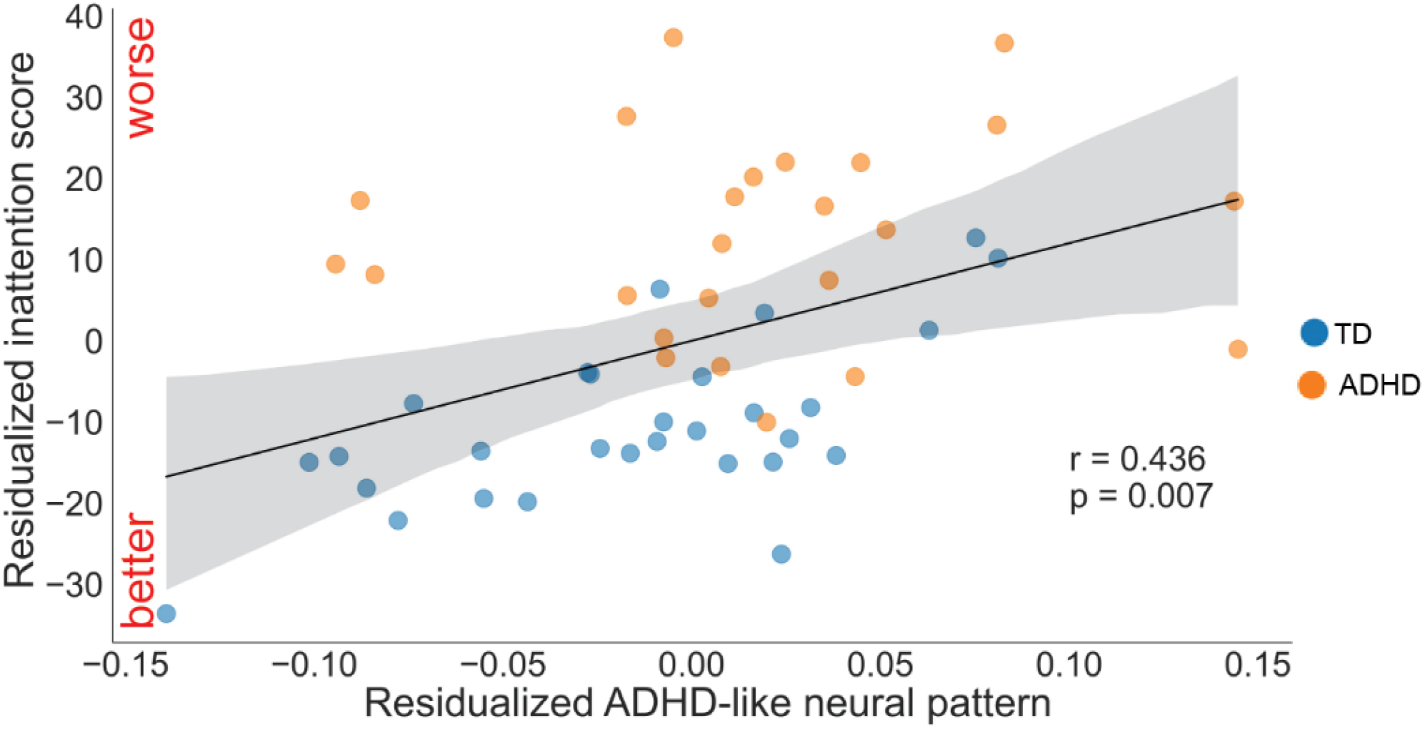
Neural variability of the rAI predicts ADHD symptom severity. ADHD-like activation pattern during proactive control in the right anterior insula (rAI) was significantly correlated with inattention scores (*r*=0.436, *p*=0.007).

### Robustness with respect to head motion

While we have meticulously excluded trials with large head motion from the single trial model (see details in Methods), head motion may remain as a potential confounding variable and bias the temporal variability and spatial stability analyses. To further test the robustness of our findings with respect to head motion, we conducted additional analyses and replicated the main findings (see **Supplementary Method, Supplementary Results and Supplementary Figure S3 – S5**).

## Discussion

We used cutting-edge analytic methodology to provide a new window into the temporal and spatial stability of neural dynamics elicited by proactive and reactive control processes in children with ADHD. We modeled single trial brain responses and utilized representational similarity analysis to relate neural coding stability to heterogeneity, temporal fluctuation in behavior, and clinical symptoms. Children with ADHD showed heightened temporal variability in task-evoked responses within key regions of the SN and FPN that are associated with behavioral variability across trials. They also demonstrated reduced spatial stability of activation patterns supporting both reactive stopping and proactive response preparation. Additionally, we observed decreased homogeneity of within-group response patterns in ADHD, indicating more variable neural recruitment strategies between children with ADHD. Importantly, indices of increased variability were further associated with more severe attention deficits and hyperactive/impulsive symptoms.

Our findings underscore fundamentally disrupted stability of neural coding across trials in ADHD leading to inefficient implementation of cognitive control mechanisms, providing a potential mechanism for inconsistent behavioral regulation. Our novel analytical approach advances knowledge of functional brain abnormalities in ADHD, moving beyond gross activation deficits toward a more precise mechanistic understanding. Our findings may help catalyze more targeted models of inhibitory control dysfunction and neurobiological heterogeneity in ADHD.

### Heightened temporal variability in ADHD

Temporal instability of cognitive and behavioral processes is increasingly recognized as a core feature of ADHD ^7,10,52,53,62^. Neural correlates of this intra-individual variability could manifest as increased fluctuation of task-evoked responses that standard averaging approaches can miss. To address this, we sought to elucidate moment-to-moment changes in inhibitory control processes by modeling single-trial variability in neural response. We expected that this novel approach would reveal instability in neural dynamics obscured in traditional group comparisons, providing a more precise understanding of cognitive control deficits in ADHD.

In the present study, we examined temporal variability of brain activity during the cued stop-signal task (CSST) using single-trial modeling. Specifically, we estimated trial-specific β time series and fitted them to a Gaussian model to derive metrics of neural variability, including standard deviation, skewness, and kurtosis. We found that children with ADHD exhibited elevated temporal variability compared to TD children. This was evidenced by increased standard deviation and kurtosis, as well as decreased skewness, of trial-wise brain responses in multiple prefrontal, parietal, and temporal cortex regions across all CSST conditions (**Figure 2)**. This suggests that children with ADHD have more variable and extreme activation levels from trial to trial than TD children.

Our results align with previous work showing more variable and noisier intrinsic brain activity in ADHD. For example, Cai et al. (2018) found more variable brain states during resting-state fMRI in children with ADHD compared to controls. Similarly, several resting-state EEG and fMRI studies have reported increased neural variability and noise in ADHD across different brain regions and frequency bands ^63–67^. Our findings extend these results by demonstrating heightened temporal variability of task-evoked neural responses in ADHD. While previous studies focused on intrinsic or resting-state brain activity, we show that the increased variability persists during cognitive demands and affects task-relevant brain regions. This task-based neural variability may have important implications for cognitive and behavioral flexibility.

Taken together, our findings provide clear evidence of heighted trial-wise neural variability in ADHD. The more temporally unstable and extreme task-related brain responses observed in ADHD likely contribute to the overall lack of consistency in performance. The noisier neural dynamics may reflect inefficiencies in cognitive control processes necessary for maintaining stable attention and motor planning over time. These results build upon a growing literature recognizing temporal variability as a core feature of ADHD and provide novel evidence linking this instability to dysfunctional inhibitory control networks. Increased neural variability may thus be a key mechanism underlying the hallmark behavioral inconsistency in ADHD, offering new insights into the neurobiology of this disorder.

### Reduced spatial stability of neural representations in ADHD

Beyond temporal variability, we also examined the spatial stability of trial-evoked brain responses during task performance. Spatial stability was defined as the similarity in activation patterns between pairs of trials within a given condition, and quantified using representational similarity analysis ^56^. RSA provides a powerful tool for investigating the distributed coding properties of cognitive control networks ^68,69^. By quantifying pattern similarity both between and within individuals across conditions, RSA allows for direct measurement of the consistency and distinctiveness of neural representations. Recent studies have begun to apply RSA to investigate interference control in the frontoparietal network ^68,70^. However, to our knowledge, no study has used trial-based RSA to examine inhibitory control in ADHD.

We computed the spatial stability of each voxel by measuring pattern similarity among neighboring voxels and used a searchlight algorithm to generate whole-brain maps of task-dependent spatial stability. Compared to controls, children with ADHD exhibited less stable activation patterns (i.e., greater spatial variability) within and across all task conditions (Uncertain Go, Certain Go, and Successful Stop, **Figure 3a**). Importantly, this reduced spatial stability was observed in key regions of the salience and frontoparietal networks, including AI, inferior frontal gyrus, middle frontal gyrus, dmPFC. These findings suggest that the salience and frontoparietal networks, critical for cognitive control, are recruited more variably from trial to trial in children with ADHD compared to TD children.

### Stability of spatial neural representations associated with reactive and proactive control

Next, we examined the spatial stability of neural representations associated with reactive and proactive control. Reactive control was assessed by contrasting Successful Stop and Uncertain Go trials: participants deployed proactive control in both these types of trials, but reactive control process was only triggered in Stop trials. Consistent with previous studies ^19,20,27^, we found that reactive control recruited key regions in the salience and frontoparietal networks (**Figure 3b**).

Crucially, TD children showed high spatial stability in widely distributed regions overlapping with the salience and frontoparietal networks during reactive control, while children with ADHD showed high spatial stability in more restricted regions, including bilateral visual cortex and right supramarginal gyrus.

Uniquely, our experimental design involved both Uncertain Go and Certain Go trials, which allowed us to examine proactive control, an aspect that has eluded previous studies and has required complex computational modeling to uncover ^15,60,71^. While some studies have suggested that proactive control recruits a highly similar brain system as reactive control^43,45,46,49–51^, the spatial stability of neural representations associated with proactive control in children with ADHD remained unknown.

In the present study, proactive control was assessed by contrasting Uncertain Go and Certain Go trials, as participants proactively slowed their responses in Uncertain Go trials. We found that TD children showed high spatial stability in the anterior insula and the inferior frontal gyrus during proactive control, but this pattern was not observed in children with ADHD. The reduced spatial stability in children with ADHD during proactive control provides novel insights into the neural mechanisms underlying the difficulties in maintaining and implementing preparatory control strategies often observed in this population ^15,72^.

Our findings suggest that while proactive and reactive control engage similar brain networks, particularly in the salience and frontoparietal regions, the consistency of recruitment within these networks differs between TD children and children with ADHD. This difference in spatial stability may contribute to a broad range of cognitive control deficits observed in ADHD. Specifically, our results indicate that the inhibitory control system was not recruited in a consistent manner from trial to trial in children with ADHD during both proactive and reactive control. The overlap in the salience and frontoparietal networks suggests that the instability of neural representations in these systems may be a common mechanism underlying deficits in both aspects of cognitive control.

In summary, our findings highlight the importance of considering the stability of neural representations associated with both proactive and reactive control in understanding the neurocognitive deficits in ADHD. The reduced consistency of recruitment within the salience and frontoparietal networks may be a key factor contributing to the cognitive control challenges faced by children with ADHD, and this may inform the development of targeted interventions aimed at improving the stability of these neural representations.

### Weak association between proactive control system and behavioral regulation in ADHD

Given the similarity in brain systems recruited by proactive and reactive control ^43,45,46,49–51^, we hypothesized that if a Go trial elicits an activation pattern resembling the typical activation pattern of reactive control, the Go trial likely involves strong proactive control, leading to elongated reaction times. This reflects the parallel neural activation patterns involved in maintaining/anticipating and executing task demands during uncertain Go and Success Stop trials. To test this hypothesis, we first created a typical brain activation pattern of reactive control by averaging trial-evoked brain responses across all Stop trials for each participant. We then developed an Inhibitory Control Pattern (ICP) index by computing the similarity between the trial-evoked response in each Go trial and the typical activation pattern of reactive control. We examined the correlation between the trial-wise ICP index and RT, with a high correlation indicating that greater spatial similarity between Go trial-evoked response and the inhibitory control pattern is associated with longer RTs. Notably, we found that the ICP index in key regions of the salience and frontoparietal network nodes tracked trial-wise RT fluctuations in TD children. However, this brain-behavior association was much weaker in children with ADHD compared to TD children (**Figure 4**).

Our findings have several important implications. First, the results align with our hypothesis that proactive control recruits a similar inhibitory control system as reactive control. Second, it suggests that spatial variability in Go trials, specifically the degree to which the trial-evoked pattern resembles activation elicited by reactive inhibitory control, encodes the amount of proactive control implemented by TD children. Third, spatial variability of trial-evoked brain responses in children with ADHD is less associated with trial-by-trial adjustment of proactive control compared to TD children.

These findings provide novel insights into the neural mechanisms underlying the behavioral regulation deficits observed in ADHD. While children with ADHD may have equivalent response slowing in Uncertain Go than Certain Go trials as TD children, children with ADHD demonstrated significant deficits in proactively adaptively their response strategy ^15^. This impairment in adaptive proactive control may contribute to the increased behavioral variability and inconsistency often observed in ADHD.

### Greater heterogeneity of trial-evoked neural response patterns in ADHD

ADHD is a highly heterogeneous disorder, with substantial variability in symptom profiles and cognitive deficits across individuals ^8,73,74^. Yet little is known about heterogeneity in task-dependent neural responses across individuals with ADHD, which may contribute to the inconsistent findings in the neuroimaging literature, as highlighted by recent studies ^74,75^. To address this gap, we first examined within-group spatial similarity during proactive and reactive control in each participant. Within-group spatial similarity was defined as the extent to which a participant’s brain activation pattern resembles the averaged activation pattern of their respective group (i.e., ADHD or TD). We hypothesized that a more homogeneous group would show greater within-group spatial similarity compared to a less homogeneous group.

Consistent with our hypothesis, we found that TD children showed greater within-group spatial similarity in a widely distributed brain system during both reactive and proactive control. In contrast, children with ADHD exhibited within-group similarity in more restricted brain regions implicated in sensorimotor processing (**Figure 5b**). This finding suggests that children with ADHD recruit more heterogeneous brain systems during task performance, which may reflect the diverse cognitive and behavioral profiles observed in this population.

To further investigate the group specificity of neural response patterns, we developed a measure of group-specific spatial similarity by subtracting between-group spatial similarity from within-group spatial similarity. Between-group spatial similarity was defined as the extent to which a participant’s brain activation pattern resembles the average activation pattern of the opposite group. Thus, the spatial similarity for a TD child measured the degree to which the child’s activation pattern is more similar to the averaged activation pattern among TD children than the averaged activation pattern among children with ADHD. Using this approach, we found that TD children showed high group specificity of spatial similarity in the inferior frontal gyrus and PPC during reactive control, and in the dmPFC and subcortical regions during proactive control (**Figure 5c**). These findings suggest that TD children recruit these regions more consistently during cognitive control tasks, whereas children with ADHD show more variable recruitment of these areas.

The heterogeneity in neural response patterns observed in children with ADHD may stem from several factors. First, the variability in symptom profiles and cognitive deficits across individuals with ADHD may be associated with distinct neural mechanisms ^8,74,76^. Second, children with ADHD may employ more diverse cognitive strategies during task performance, leading to greater variability in brain activation patterns ^8,77^. Finally, the inherent instability of neural dynamics in ADHD, as demonstrated by our findings of increased temporal variability and reduced spatial stability, may also contribute to the heterogeneity of neural response patterns.

Taken together, our findings reveal that children with ADHD exhibit more heterogeneous neural response patterns during cognitive control tasks compared to TD children. This neural heterogeneity may reflect the diverse clinical presentations and cognitive profiles observed in ADHD. Our approach of examining within-group and between-group spatial similarity provides a novel framework for investigating the neural basis of symptom variability in ADHD and other neuropsychiatric disorders.

### Temporal variability and spatial pattern similarity predict ADHD clinical symptoms

Having established that children with ADHD exhibit increased temporal variability and reduced spatial stability of neural responses during cognitive control tasks, we next investigated whether these neural measures could predict individual differences in ADHD clinical symptoms. We focused on two core symptom domains: inattention and hyperactivity/impulsivity.

We found that higher temporal variability of trial-evoked neural responses in one key region of the salience and frontoparietal networks, the right PPC, was associated with more severe inattention symptoms. This finding aligns with previous research suggesting a potential association between stimulation of the right PPC region and improvements in attentional control during task performance ^78^. It suggests that the increased neural variability observed in children with ADHD is not merely a reflection of general neurodevelopmental differences but is directly related to the severity of clinical impairments. The link between temporal variability and symptom severity may stem from the role of these networks in maintaining stable goal-directed behavior and suppressing irrelevant distractors ^27,28,69,79–84^. Increased variability in these networks may lead to more frequent lapses of attention and difficulty inhibiting impulsive responses, thereby exacerbating ADHD symptoms.

Furthermore, we found that children who exhibited more ADHD-like spatial activation patterns during proactive control had more severe inattention symptoms (**Figure 6**). This finding aligns with our previous work showing that children with less adult-like activation patterns during inhibitory control tasks had weaker overall inhibitory control function ^85^. However, our previous study focused on averaged activation patterns across trials, potentially obscuring trial-by-trial variability in neural responses ^85^. The current findings extend our previous work by demonstrating that the spatial stability of trial-evoked brain responses, rather than just the average activation pattern, is predictive of ADHD symptom severity. This novel metric of neural stability may provide a more robust measure of the neural mechanisms underlying ADHD symptoms and may serve as a potential biomarker for the disorder.

The association between more ADHD-like spatial activation patterns and more severe inattention symptoms suggests that children with ADHD who can maintain more stable and typical neural representations during cognitive control tasks may be better able to regulate their attention and behavior. This finding highlights the importance of considering not only the magnitude of neural activation but also the consistency and typicality of neural response patterns when investigating the neural basis of ADHD symptoms.

Our results have important implications for the development of novel interventions targeting the neural mechanisms underlying ADHD symptoms. For example, interventions that aim to enhance the stability of neural responses in the salience and frontoparietal networks, such as neurofeedback training ^86–89^, may help to reduce symptom severity in children with ADHD. Additionally, our findings suggest that measuring the spatial stability of trial-evoked neural responses could serve as a useful metric for tracking treatment response and predicting long-term outcomes in ADHD.

## Conclusion

We employed cutting-edge neuroimaging techniques and innovative analytical approaches to investigate the neural mechanisms underlying dynamic, proactive, and reactive control processes in children with ADHD. By examining the temporal variability and spatial stability of trial-evoked brain responses, we provide novel insights into the neural underpinnings of cognitive control deficits in ADHD, which have been poorly understood due to the limitations of traditional neuroimaging methods.

We demonstrate that children with ADHD exhibited increased temporal variability and weakened spatial stability of trial-evoked brain responses in the salience and frontoparietal networks compared to typically developing children. Furthermore, TD children showed highly similar spatial patterns of neural activity, whereas children with ADHD demonstrated more heterogeneous and idiosyncratic patterns. Crucially, the temporal variability and spatial pattern similarity of neural responses were functionally and clinically relevant. In TD children, spatial stability in the salience and frontoparietal networks tracked trial-by-trial fluctuations in response time, but this effect was much weaker in children with ADHD. Moreover, neural variability measures were related to the severity of core ADHD symptoms, such as inattention and hyperactivity/impulsivity.

Our findings provide novel insights into the dynamic and heterogeneous neural abnormalities underlying cognitive control deficits and symptom variability in childhood ADHD. These results highlight the potential of neural variability measures as biomarkers for ADHD and as targets for innovative interventions aimed at enhancing the stability and consistency of neural activity patterns. Our study demonstrates the utility of advanced neuroimaging approaches in uncovering subtle and dynamic neural mechanisms in ADHD and other neurodevelopmental disorders, paving the way for personalized interventions and improved outcomes.

## Methods

### Participants

One hundred and seven children (9–12 years old) were recruited from the local community. Informed consent was obtained from legal guardians of the children and the study was approved by the Institutional Review Board of Stanford University. Ninety-eight children completed two runs of CSST in the scanner. Thirty-seven children were excluded in the analysis because of missing data, image artifacts, excessive head motion (criteria: mean frame displacement (FD) was larger than 0.25mm, max frame displacement was larger than 5mm), and behavioral outliers (criteria: mean Go trials accuracy was less than 75%, stop accuracy was lower than 25% or higher than 75%) ^90^. The final dataset included 26 children with ADHD (10Lfemale, 16Lmale) and 35 TD children (13Lfemale, 22Lmale).

### Clinical and neuropsychological assessments

Children and their guardians completed a clinical and neuropsychological assessment session. ADHD diagnosis was informed by the children’s guardians and further confirmed using the Conners 3^rd^ Edition. ADHD with conduct disorder and oppositional defiant disorder were not excluded because of their high comorbidity rates ^91^. Additional enrollment criterion for both children with ADHD and TD children included no history of claustrophobia, head injury, serious neurological or medical illness, autism, psychosis, mania/bipolar, major depression, learning disability, substance abuse, sensory impairment such as vision or hearing loss, birth weight <2000 g and/or gestational ages of <34 weeks. All children were right-handed with an IQ >80.

Inattention and hyperactivity/impulsivity symptoms were assessed using The Conners’ Rating Scale (Parent). Participants who were under stimulant treatment had gone through a washout period of at least 5 half-lives of the medicine before testing.

### Cued stop-signal task (CSST)

The CSST was modified from the standard stop-signal task in order to dissociate proactive control from reactive control processes (**Figure 1a**) ^43,46^. Each trial began with a white or green cross (Cue) in the center of the screen for 200 milliseconds, followed by a green arrow (go signal). Participants were demanded to make an accurate and speedy button press in response to the pointing direction of the green arrow. Occasionally, the green arrow turned to red (stop signal), and participants had to withhold their responses when the color changed. The color of the cue indicated the probability of the stop signal in the coming trial. A green cross indicates that no stop signal would occur after the coming go signal, which is defined as **Certain Go** trial. A white cross indicates that a stop signal may occur after the coming go signal (33% chance), which is defined as **Uncertain Go** trial. If a stop signal is presented, the trial is defined as **Stop** trial. The stop-signal delay (SSD) was initialized at 200ms and adjusted in a staircase fashion. If the participant successfully canceled a prepotent response, the SSD increased by 50ms in the next Stop trial. If the participant failed in stopping, the SSD decreased by 50ms in the next Stop trial. Participants completed two runs of the CSST in the scanner, and each run included 32 Certain Go trials, 32 Uncertain Go trials, and 16 Stop trials with jittered inter-trial intervals (ITIs) between 1 and 4 seconds.

### Behavioral measures

Reactive control was measured by SSRT. First, we confirmed that behavioral data did not violate the main assumption of the Race Model, that the mean RT in Unsuccessful Stop (US) trials should be shorter than the mean RT in Go trials. Then, SSRT was computed using the integration method based on the Race model ^17,90^: SSRT = T−mean SSD, where T is the cutoff point where the integral of the observed distribution of Go RT in the SST (or Uncertain Go RT in the CSST) equals the probability of unsuccessful stopping. Proactive control was measured by response slowing modulated by task cues, i.e. Uncertain Go RT minus Certain Go RT.

### Neuroimaging data acquisition

Imaging data were acquired on a 3.0 T GE Signa scanner using a 32-channel head coil at the Richard M Lucas Center for Imaging at Stanford University. Functional images of 42 axial slices were acquired using the multiband gradient-echo planar imaging with the following parameters: TR=490ms; TE=30ms; flip angle=45°, FOV=22.2cm, matrix=74x74 and in-plane resolution=3 mm. A high-order shimming method was used prior to data acquisition to reduce blurring and signal loss arising from field inhomogeneity. High-resolution T1-weighted images were acquired using a spoiled-gradient-recalled inversion recovery three-dimensional (3D) MRI sequence with the following parameters: TR=8.4ms, TE=1.8ms, flip angle=15°, FOV = 22cm, matrix=256x192.

### fMRI data pre-processing analysis

Image pre-processing and statistical analysis were performed using SPM12 (https://www.fil.ion.ucl.ac.uk/spm/software/spm12). The first 12 volumes before the task were discarded to allow for T1 equilibrium. The remaining images were then realigned to correct for head movements and underwent slice-timing correction. EPI images were registered to the MNI standard space. Data were spatially smoothed using a 2 mm FWHM Gaussian kernel as recent studies suggested that minimal spatial smoothing could enhance signal-to-noise ratio while preserving distributed pattern information ^92^.

### General linear model (GLM)

Conventional GLM was conducted to estimate trial averaged neural responses elicited by task conditions in the CSST, including Certain Go, Uncertain Go, Successful Stop (SuccStop), Unsuccessful Stop (UnsuccStop) and Go error. Six motion parameters were entered as covariates of no interest.

### Single-trial GLM

The GLMs were performed separately to estimate the activation pattern for each trial using a Least Square–Separate (LS-S) approach ^93,94^, in which the trial of interest was modelled as one regressor, with all other trials modelled as separate regressors. Specifically, each single-trial GLM included five regressors: (1) the single trial of interest from one condition, for instance one Successful Stop trial (SS); (2) all other SS trials; (3) all the Uncertain Go trials; (4) all the Certain Go trials; (5) all the US trials. Six head motion parameters were included as regressors of no interest. Each event was modelled by defining the time of stimulus onset as a stick function and convolving this stick function with a canonical hemodynamic response function. Temporal autocorrelations were modelled with the FAST model ^95^. This voxel-wise GLM was used to compute the activation associated with each trial. The β-map for each trial was used for the main analyses, including temporal variability and spatial stability analyses. Crucially, in order to minimize the impact of head movements, we removed trials in which head motion exceeded 0.5mm framewise displacement (FD) within the 4 to 8 seconds after stimulus onset, a method widely used in previous studies ^96–98^. These exclusions accounted for 0.28% and 1.52% of the volume for TD and ADHD groups, respectively.

### Temporal variability analysis

We first assessed the normality of the distribution of trial-evoked brain responses (β time series) (**Supplementary Methods and Table S5**). Then, the β time series were fitted with the Gaussian Model per voxel and per condition, and four parameters were derived, including mean, standard deviation, skewness, and kurtosis. Given the limited number of SS and US trials, we merged these two types of trials into one ‘Stop’ condition in this analysis. FSL’s Randomise procedure was implemented to determine statistical significance with 5,000 permutations and results were thresholded at *p*<0.05 TFCE corrected (Threshold-Free Cluster Enhancement) ^99–101^. Of note, the significance map generated from FSL’s Randomise represents values of 1 minus p.

### Spatial stability analysis

We utilized representational similarity analysis (RSA) employing both searchlight and regions of interest (ROI) methods ^56,58^ to examine spatial stability during task performance. For searchlight analysis, β-maps were obtained from a cubic region of interest containing 125 surrounding voxels across each participant’s whole brain (as in Viganò and Piazza ^102,^Xu, et al. ^103,^Gao, et al. ^104,^Zheng, et al. ^105^). To calculate pattern similarity, we correlated activity vectors of any given pair of trials using Pearson correlation. Subsequently, we transformed these similarity scores into Fisher’s z-scores and compared them between conditions to quantify proactive and reactive control. ROI-based analyses were conducted similarly, except that the Pearson correlation was computed using voxels specifically selected from the chosen ROI. Of note, we excluded any pairs presented in the same run from the calculation of pattern similarity to avoid any autocorrelation issues.

Spatial stability of trial-evoked brain responses was calculated in Uncertain Go, Certain Go, and Successful Stop trials for each participant, then between-group comparisons were conducted to examine whether the spatial stability was influenced by the disorder. Proactive control was quantified by the contrast of within-condition pattern similarity between Uncertain Go and Certain Go trials, as the sole distinction between these two conditions lies in the inclusion of a proactive control component in Uncertain Go but not in Certain Go trials. Similarly, reactive control was quantified by contrasting Successful Stop trials with Uncertain Go trials, given that only the former incorporates a reactive component (stopping). The contrasting maps were used for group-level analysis and between-group comparison. Statistical significance was tested using FSL’s ‘randomize’ like previously mentioned.

### Pattern similarity of inhibitory control tracks RT fluctuation

First, we created an activation pattern template of inhibitory control by averaging trial-evoked activation patterns (i.e., single trial β maps) across all successful stop trials for each participant. Then, we computed the similarity of activation patterns between the inhibitory control template and each go trial, resulting a time series of correlation coefficients along all the go trials. Next, we correlated the resulting time series of pattern similarity with the RT across go trials, using the searchlight approach. The correlation coefficient maps were z transformed and subsequently used for group-level comparison. Statistical significance was tested using FSL’s ‘randomize’ like previously mentioned.

### Group-specific inter-subject similarity analysis

We developed a group-specific inter-subject similarity index to characterize the extent to which a participant’s brain activation pattern is similar to the rest of the participants from the same group in relative to the participants from the other group ^106^. For each participant, we averaged all other participants’ β-maps from the same group (ADHD or TD) for each condition. Each participant’s β map was then correlated with the average maps either from their own group or from the other group, resulting within-group inter-subject similarity map and between-group inter-subject similarity map, respectively. The group-specificity inter-subject similarity index was calculated by subtracting the between-group inter-subject similarity map from the within-group inter-subject similarity map. We compared these pattern similarity measures between conditions, similar to a within-participants analysis, to investigate whether children with and without ADHD exhibited group-specific neural activity patterns related to proactive and reactive control. This analysis was conducted in a whole-brain searchlight approach. The significance test was conducted using permutation test implemented in FSL randomize, as previously mentioned.

### Temporal variability and spatial pattern similarity in association with clinical symptoms

To investigate whether temporal variability and spatial pattern similarity could explain individual differences in core symptoms of ADHD, we conducted ROI-based correlation analysis. Our focus was on the core regions implicated in cognitive control (see the below Regions of Interest in the Method part). Temporal variability of brain responses was extracted from independent defined ROIs, and clinical symptoms, including hyperactivity and inattention scores, were assessed using Conners’ Rating Scale. Pearson’s correlation was employed to evaluate their relationship. Regarding spatial pattern similarity and its link to clinical symptoms, our interest lies in examining the presence of ADHD-like neural activity patterns supporting proactive and reactive control. Thus, we conducted ROI-based analysis to investigate whether the similarity to the averaged spatial patterns of the TD group is was associated with clinical symptoms of ADHD.

### Regions of interest (ROI) Analysis

To examine the robustness of our findings and further confirm children with ADHD demonstrated increased temporal variability and decreased spatial stability in the salience and frontal-parietal network compared to TD children. We conducted additional analyses focusing on the core cognitive control regions. Task-relevant brain regions involved in cognitive control were defined using functional clusters from an independent study, where brain networks were derived using independent component analysis on resting-state fMRI data ^107^. Specifically, we selected the right anterior insula (rAI) and dorsal medial prefrontal cortex (dmPFC) from the salience network, and the right dorsolateral prefrontal cortex (rdlPFC) and right posterior parietal cortex (rPPC) from the right executive control network.

## Data availability

Original data reported in this study is available through request.

## Code availability

Functional MRI data preprocessing and statistical analyses were performed on the SPM12 and FSL 6, and Matlab 2020.

## Acknowledgements

This research was supported by National Institutes of Health MH105625 (WC), MH124816 (WC) MH121069 (VM), EB022907(VM) and NS086085 (VM), NARSAD Young Investigator Award (WC), Stanford Maternal and Child Health Research Institute Grant (WC), and Stanford University Department of Psychiatry Innovator Grant (WC). We thank Rachel Rehert and Ahmad Belai AI-Zughoul for their assistance with data collection.

## Author contributions

Study design: W.C., V.M.; Data collection: W.C., K.D., S.W.; Data Analysis: Z.G., W.C., L.Z.;

Manuscript Drafting: Z.G., W.C.; Manuscript Editing: Z.G., S.W., S.H., V.M., W.C.

## Notes

### Competing Interest Statement

The authors have declared no competing interest.

