## supplementary materials for "Reduced temporal and spatial stability of neural activity patterns predict cognitive control deficits in children with ADHD"

**Supplementary Methods**

**Normality of distribution of trial-evoked neural responses**

Prior to assessing the temporal variability of trial-evoked neural responses, we tested determine whether it is appropriate to fit trial-evoked neural responses in the Gaussian model. The test specifically targeted 17 ROIs of the inhibitory control system, as identified by an independent meta-analysis study (Cai et al., 2019). The β time series for each trial type (e.g., Uncertain Go, Certain Go, and Stop) were extracted from the peak voxel within each ROI for each participant, and subsequently fitted with a normal distribution. The assessment of normality was performed using the 'Chi2gof' function available in Matlab 2020. The proportion of non-normally distributed data was computed for each ROI and condition.

**Consistency of recruitment within the salience and frontoparietal network between proactive and reactive control**

Our study found that both proactive and reactive control engaged similar brain networks, particularly within the salience and frontoparietal regions. However, there are notable differences in the consistency of recruitment within these networks between typically developing (TD) children and children with ADHD. To validate this observation, we investigated the spatial consistency of recruitment of proactive and reactive control within the salience and frontoparietal network for each group. Reactive control was identified using the conventional univariate contrast between Successful Stop trials and Uncertain Go trials, while proactive control was delineated by the contrast between Uncertain Go and Certain Go trials. *Pearson’s* correlation was used to assess the consistency of recruitment between these two control processes by correlating the statistic maps within the salience and frontoparietal network.

**Controlling the potential confounding factor of head motion**

Although we excluded trials with large head motion from the single trial models, head motion may still remain as a potential confounding variable. To further mitigate this concern, we repeated the main analyses while controlling the remaining head motion effect (see **Supplementary Figure S3 – S5**). Specifically, mean frame displacement for each participant was included as covariate. We demonstrated that the main findings are robust against head motion effect.

**Supplementary Results**

**Normality of distribution of trial-evoked neural responses**

We found that trial-evoked neural responses in our study fit a normal distribution for most of our participants, ranging between 82% and 97% (see **Supplementary Table S2**). There were no significant between-group differences in non-normality (all ps > 0.2).

**Consistency of recruitment within the salience and frontoparietal network between proactive and reactive control**

In comparison to children with ADHD (*r*=0.05), TD children (*r*=0.17) exhibited significantly higher consistency in recruitment of the SN and FPN between proactive and reactive control (*z*=14.94, *p*<0.001).

**Supplementary Figures**

**Figure S1.** Children with ADHD showed lower skewness of trial-evoked brain responses in comparison to TD children (*p*<0.05, corrected).
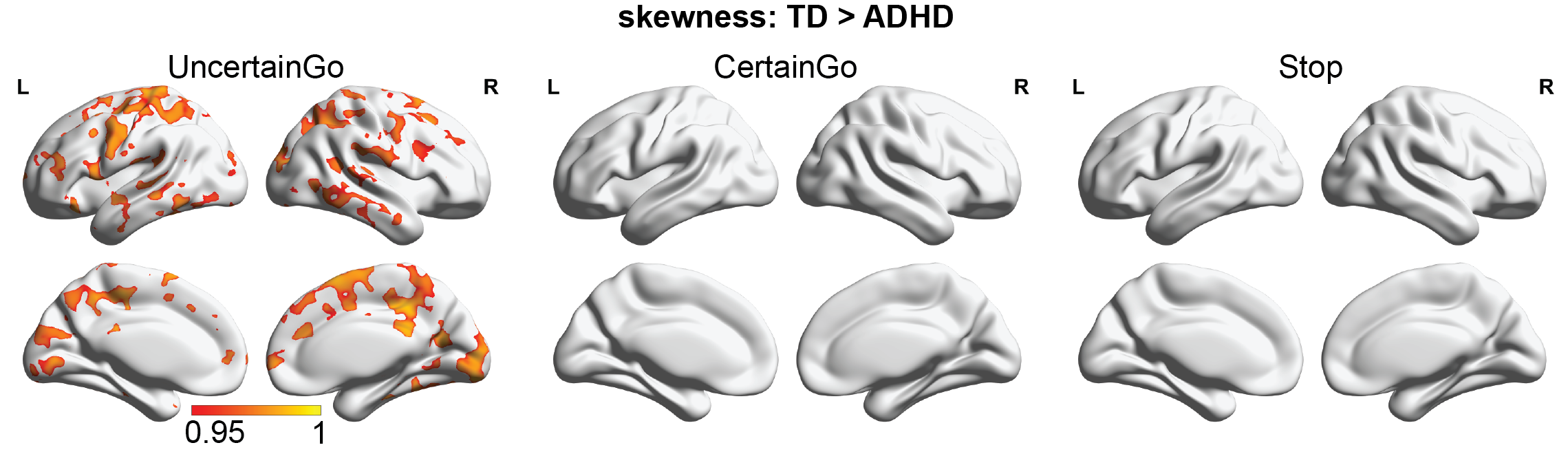

**Figure S2.** Children with ADHD showed less pattern similarity supporting proactive and reactive control in comparison to TD children when using a less stringent threshold (*p*<0.01, uncorrected).

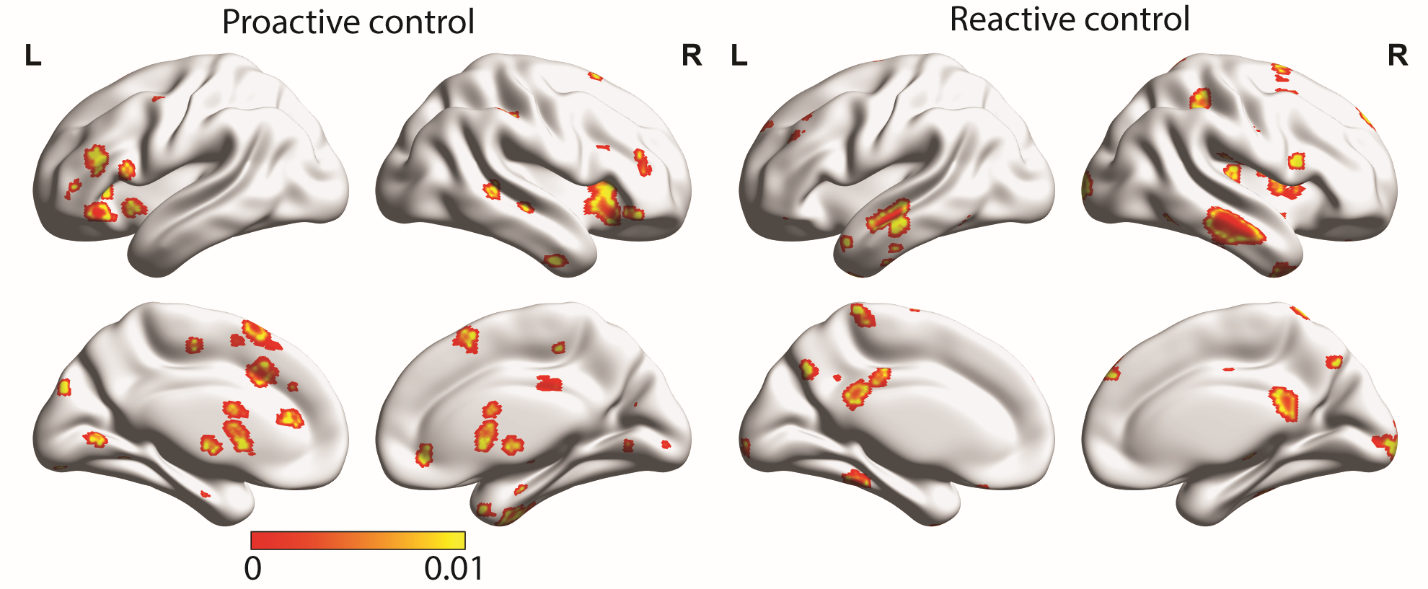

**Figure S3.** Children with ADHD showed significantly higher standard deviations of neural activity compared to TD children across all three trial types. Children with ADHD also exhibited significantly greater kurtosis during Uncertain and Certain Go trials compared to TD children. TD children demonstrated greater skewness of neural activity during Uncertain Go trials when compared to children with ADHD. All results underwent TFCE correction, *p*<0.05 (of note, the significance map represents values of 1-p). Analyses were conducted with additional head motion control.

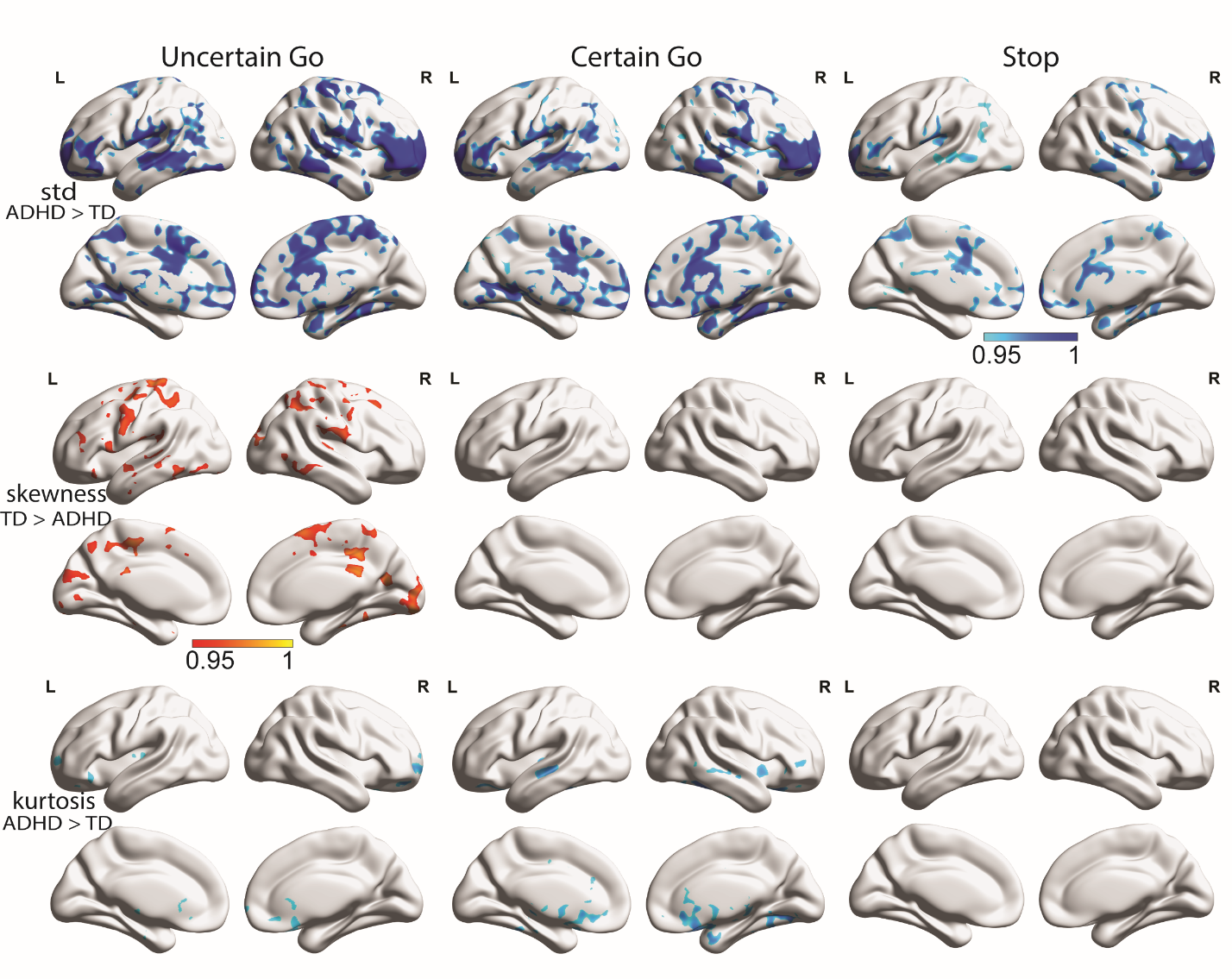

**Figure S4.** In comparison to children with ADHD, TD children demonstrated significant greater spatial stability of neural activity patterns across three trial types after controlling head motion (TFCE, *p*<0.05, the significance map represents values of 1 minus p).

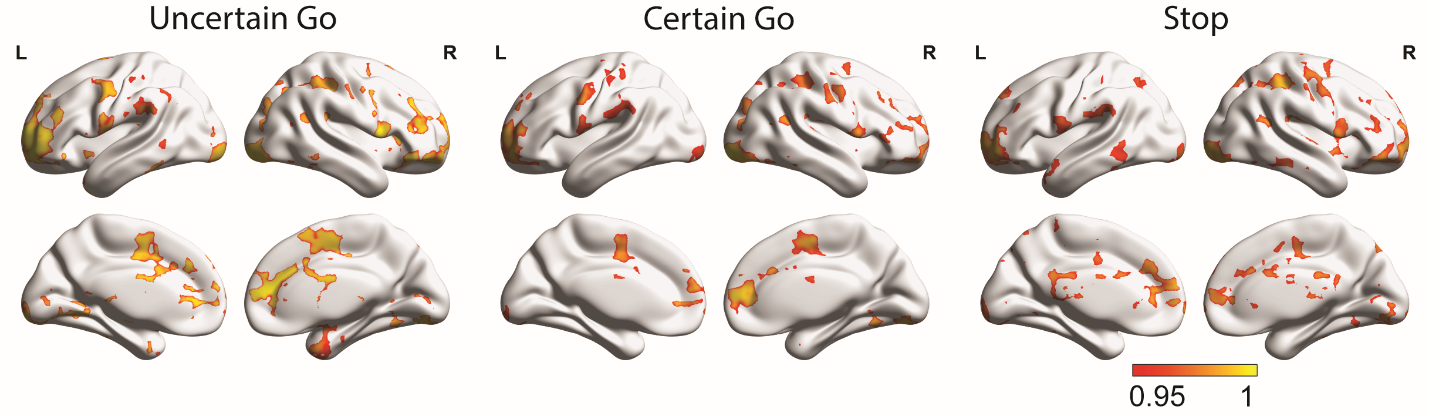

**Figure S5.** Weak association between trial-evoked inhibition-alike brain response and RT fluctuation in children with ADHD during Go trials (TFCE, *p*<0.05, the significance map represents values of 1 minus p). Analyses were conducted with additional head motion control.

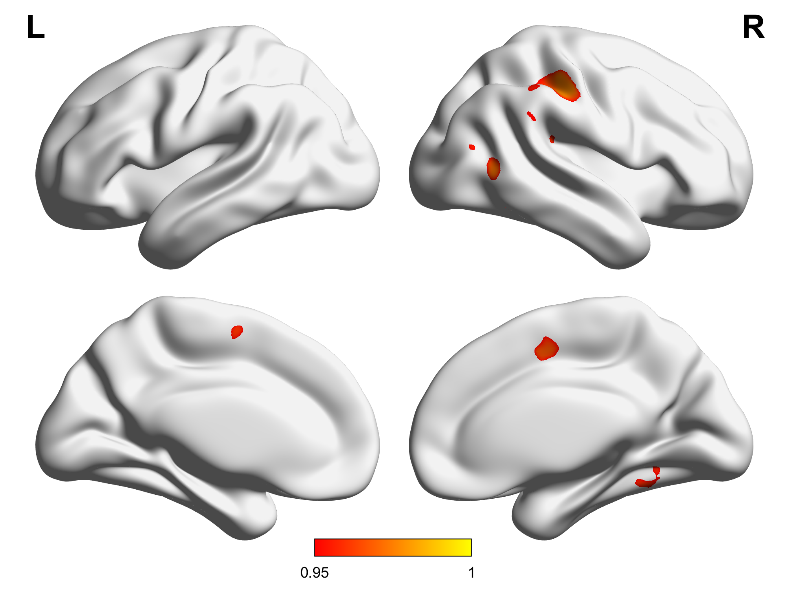

**Supplementary Tables**

**Table S1.** Demographic and behavioral statistics of behavioral-only sample, including all participants who meet the behavioral criteria regardless of their head motion.

| Demographics | TD Controls (N=37) | ADHD (N = 50) | p-value |
| --- | --- | --- | --- |
| Age (SD) | 10.46 (1.07) | 10.66 (1.20) | 0.406 |
| Gender (Male/Female) | 23/14 | 35/15 | 0.443 |
| IQ (SD) | 113.649 (13.292) | 103.041 (14.177) | p<0.001 |
| Inattention score | 49.55 | 79.85 | p<0.001 |
| Hyperactivity score | 49.32 | 79.09 | p<0.001 |
| Cued SST Performance |  |  |  |
| Certain Go Accuracy (%) | 92.5 | 90.8 | 0.234 |
| Uncertain Go Accuracy (%) | 94.8 | 91.4 | 0.017 |
| Stop Accuracy (%) | 52.2 | 52.4 | 0.898 |
| SSRT (ms) | 296.6 | 337.4 | 0.016 |
| RT slowing (ms) | 20.4 | 23.8 | 0.705 |
| Certain Go RT (ms) | 483.4 | 534.5 | 0.004 |
| Uncertain Go RT (ms) | 507.1 | 559.6 | 0.002 |
| ex-Gaussian modelling |  |  |  |
| Certain Go RT (mu) | 395.7 | 423.3 | 0.009 |
| Certain Go RT (sigma) | 109.8 | 134.9 | 0.206 |
| Certain Go RT (tau) | 87.7 | 112.2 | 0.021 |
| Uncertain Go RT (mu) | 418.4 | 438.5 | 0.055 |
| Uncertain Go RT (sigma) | 50.2 | 50.0 | 0.968 |
| Uncertain Go RT (tau) | 88.7 | 121.1 | 0.002 |

A chi-square test was performed to assess the presence of gender differences between children with ADHD and typically developing (TD) children. For all other between-group comparisons, two-tailed t-tests were utilized.

**Table S2.** The percentage of participants whose trial-evoked neural responses rejected the null hypothesis of normal distribution.

| ROI name | Uncertain Go | Certain Go | Stop |
| --- | --- | --- | --- |
| rAI | 0.15 | 0.1 | 0.11 |
| rSFG | 0.11 | 0.05 | 0.11 |
| rFP | 0.07 | 0.13 | 0.05 |
| rSPL | 0.07 | 0.11 | 0.05 |
| rSMG | 0.1 | 0.1 | 0.03 |
| lAI | 0.1 | 0.18 | 0.08 |
| rSTG | 0.07 | 0.15 | 0.05 |
| rThalamus | 0.17 | 0.15 | 0.13 |
| lSMG (posterior) | 0.15 | 0.1 | 0.07 |
| rMFG | 0.07 | 0.08 | 0.1 |
| lSMG (anterior) | 0.07 | 0.11 | 0.07 |
| Precuneus | 0.05 | 0.07 | 0.07 |
| lSPL | 0.05 | 0.08 | 0.11 |
| lSFG | 0.07 | 0.07 | 0.11 |
| lMFG | 0.13 | 0.23 | 0.1 |
| lOG | 0.15 | 0.11 | 0.1 |
| rSFG/Precentral | 0.13 | 0.1 | 0.05 |
| Averaged | 0.1 | 0.11 | 0.08 |

rAI, right anterior insula; rSFG, right superior frontal gyrus; rFP, right frontal pole; rSPL, right superior parietal lobe; rSMG, right supramarginal gyrus; lAI, left anterior insula; rSTG, right superior temporal gyrus; rThalamus, right thalamus; lSMG, left supramarginal gyrus; lSPL, left superior parietal lobe; lSFG, left superior frontal gyrus; IOG, inferior occipital gyrus
